# CBMOS: a GPU-enabled Python framework for the numerical study of center-based models

**DOI:** 10.1101/2021.05.06.442893

**Authors:** Sonja Mathias, Adrien Coulier, Andreas Hellander

## Abstract

Cell-based models are becoming increasingly popular for applications in developmental biology. However, the impact of numerical choices on the accuracy and efficiency of the simulation of these models is rarely meticulously tested. We present CBMOS, a Python framework for the simulation of the center-based or cell-centered model. Contrary to other implementations, CBMOS’ focus is on facilitating numerical study of center-based models by providing access to multiple ODE solvers and force functions through a flexible, user-friendly API. We show-case its potential by evaluating the use of the backward Euler method for calculating the trajectories of two-dimensional cell populations. We confirm that although for moderate accuracy levels the backward Euler method allows for larger time step sizes than the commonly used forward Euler method, its additional computational cost due to being an implicit method prohibits its use for practical test cases.

CBMOS is available on GitHub^1^ and PyPI under an MIT license. It allows for fast prototyping on a CPU for small systems through the use of NumPy. Using CuPy on a GPU, cell populations of up to 10,000 cells can be simulated within a few seconds. As such, we hope it can also be of use to modelers interested in testing preliminary hypotheses before committing to more complex center-based model frameworks.

**AMS subject classification:** 65Z05, 92C15, 92-10

## 1 Introduction

Cell-based models offer the possibility to study questions relating cellular to population level behavior by explicitly representing each individual cell and its mechanical interaction with its neighbors [1]. As such, they are becoming increasingly popular for applications in developmental [2, 3] and cancer biology [4, 5].

There exist a multitude of cell-based models which can be categorized as either *on-* or *off-lattice* models. On-lattice models such as cellular automata [6] or the cellular potts model [7] restrict the movement of cells in space to a fixed grid. Off-lattice models, on the other hand, track the movement of particles in continuous space. Depending on the resolution of the shape of the cells, the particles can represent cell midpoints (center-based or cell-centered model [8, 9]), cell membrane junctions (vertex-based model [10]) or even individual cell parts, such as in the immersed boundary method [11] or the subcellular element model [12]. Center-based models represent cells as overlapping spheres or as Voronoi polyhedra defined by the topology of their midpoints [9, 13]. Cells then interact mechanically with their neighbors according to pairwise forces, in an analogy to them being connected by springs. The exact definition of the cell’s neighborhood depends on the method, where the most common ones are either solely distance-based or restrict interaction to neighboring Voronoi polyhedra. Pairs of cells then attract or repel each other depending on the distance between them. Currently available software for the general simulation of center-based models include the open-source frameworks *Chaste* [14], *MecaGen* [15], *PhysiCell* [16], and *ya* ‖ *a* [17], as well as the closed-source packages *CellSys* [18], *EPISIM* [19] and *Biocellion* [20].

The robustness of biological conclusions to both basic model assumptions and numerical issues is of the utmost importance in order to build upon predictions and understanding gained from simulations. With the increased usage of cell-based models there is an increased need for the numerical study of these models for several reasons: (i) In the absence of exact data on intercellular forces in cell populations such as tissues, models abstract how cells mechanically interact with their immediate neighbors in different ways, e.g. with different types of *ad-hoc* pairwise interaction forces. Changing the exact mathematical definition of these force functions should — in the best case — have no impact on the behavior at the population level. Modellers should take care to confirm this expectation, or at the very least, study how their model conclusions are affected by these changes. (ii) The simulation of these models relies on the numerical solving of update equations for different model components, e.g. the movement of particles such as the cell midpoint coordinates. In general, it is not straightforward to know whether more complex numerical methods — e.g. a higher order method or an implicit method — are beneficial over more simpler ones in terms of computational cost necessary to achieve a needed level of accuracy. (iii) Careful consideration needs to be given to the choice of purely numerical parameters, such as time step length or number of iterations, as these have been shown to affect model conclusions [21].

Several publications have studied these three points in the context of center-based models. Such publications include the work of Pathmanathan et al. [22], where they compared the bulk mechanical properties of a non-proliferating tissue simulated using different physics-based forces, and the work by K. Atwell [23] which investigated both the use of different forces and of different numerical solvers for the simulation of a tumor growth type experiment. In particular, the latter compared a fourth-order explicit Runge-Kutta method and two implicit methods to the commonly used first-order explicit forward Euler method in terms of accuracy and run times, as well as proposing a simple adaptive mechanism to ensure that the time step size is chosen small enough to not violate a given absolute movement threshold. Additionally, in [13] Osborne et al. compared five different cell-based models — amongst others the center-based model — with respect to their underlying model assumptions, implementation details and applicability to different common biological problems. All of the above studies have been performed using the Chaste simulation framework. Furthermore, as part of the supplementary information of the publication announcing the PhysiCell code [16], Ghaffarizadeh et al. ran convergence studies for the second-order explicit Adams-Bashforth method (used by PhysiCell for updating the cell positions) for a two-cell test case and a compressed spheroid population.

In a previous study [24] we explored the question of how the formulation of the force function governing the pairwise interaction forces affects the numerical properties of the two-dimensional center-based model when used in combination with different first and second-order explicit numerical methods. We showed that, for the simulation to remain physically correct, the size of the time step must be tailored to the choice of force function. Moreover, choosing the time step size too large for a given force function/solver combination led to geometrical differences at the population level, with the different force functions exhibiting varying sensitivity to this issue. These findings illustrate the importance of ensuring that model conclusions are independent of numerical choices.

With the exception of the last one, all of the above studies have been performed within a typical feature-rich modeling software written primarily to study biological problems within the context of a specific modeling problem. These frameworks typically only provide one type of force function and one solver (note that Chaste provides general interfaces for both which the user could extend [14]). As a complementary approach, there is value in studying these issues in a more general setting in order to inform modelers on how the combination of different basic model assumptions can affect typical population level behavior and how to avoid common pitfalls with respect to numerical parameters.

To this end, we have written CBMOS, a framework designed explicitly for the numerical study of center-based models. Our code is a Python implementation making it easily accessible for novice and experienced programmers alike, while internally relying heavily on NumPy’s vectorized routines [25] for performance. Through the optional use of the CuPy library [26], it enables transfer of the calculation of the pairwise cellular forces to a graphical processing unit (GPU) if available, thus allowing for the simulation of cell population sizes of up to 10,000 cells in a few seconds.

CBMOS is publicly available on GitHub^2^ along with all Jupyter notebooks that were used to generate the figures in this publication. Additionally, it is available on the Python Package Index (PyPI) and can be installed via pip by running pip install cbmos. It has minimal requirements consisting mainly of the scientific Python stack NumPy, SciPy and Matplotlib, along with the optional requirement of CuPy if execution on the GPU is desired. Documentation can be found on the project’s GitHub page^3^. Users interested at trying out CBMOS without installing it on their system, can also choose to run it through Google Colaboratory (or Google Colab for short) [27, 28]. Google Colab offers a Python computing environment based on Jupyter notebooks with access to GPU hardware. It runs entirely in the cloud and is accessible through a web browser^4^. Most conveniently, it also features the possibility to directly run Jupyter notebooks hosted on GitHub, such as those that can be found in the CBMOS directory, enabling any potential user to quickly get started with CBMOS.

Our article is organized as follows. Section 2 describes the center-based model and the design of our code to facilitate numerical exploration, along with implementation aspects and how users can install CBMOS. In Section 3 we illustrate the performance gained by transferring the main bulk of the calculations to the GPU and probe the range of system sizes CBMOS is able to simulate efficiently. Additionally, to show-case the potential of our framework, we conduct an in-depth numerical study into the efficiency of the backward Euler method when used to calculate cell trajectories for two-dimensional populations. Lastly, we discuss the advantages and limitations of our code in Section 4.

## 2 The CBMOS software framework

In this section we briefly state the mathematical description of the center-based model as it is implemented in CBMOS and explain its design using a minimal working example. Moreover, we describe implementation aspects such as the array programming paradigm we use, as well as which parts of the code are extended to the GPU. Finally, we describe how interested users can install it or quickly set it up on Google Colab.

### 2.1 Mathematical description of the center-based model

The center-based model implemented in CBMOS tracks the movements of the midpoint coordinates **x** of a population of cells over time. Individual cells are implicitly represented as circles (in two dimensions) or spheres (in three dimensions) with a fixed radius *R*. They are assumed to interact mechanically according to pairwise interactions with their neighbors within a certain maximum interaction distance *r*_*A*_, with the magnitude of their force interaction depending only on their distance. If two neighboring cells are located closer than some rest length *s*, they exert repulsive forces on each other to eliminate their overlap. If they are placed exactly the rest length *s* apart, they exert no forces on each other. (As a default, CBMOS uses a rest length of *s* = 1.0 cell diameter.) Additionally, if located in close proximity, but not overlapping (i.e. at a distance larger than the rest length *s* but smaller than the maximum interaction distance *r*_*A*_), they exert adhesive forces pulling them closer. Specific pairwise force functions implementing such behavior include the cubic force implemented in MecaGen,

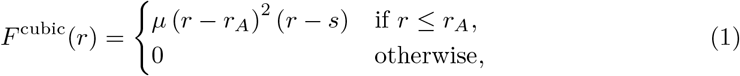

the piecewise quadratic (PWQ) force used in PhysiCell,

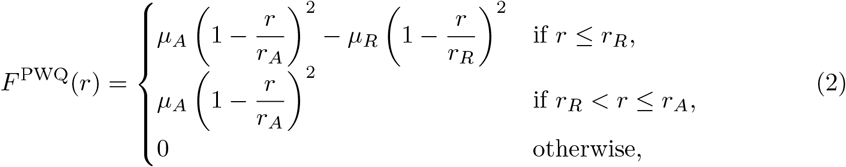

and the generalized linear spring (GLS) force used in Chaste,

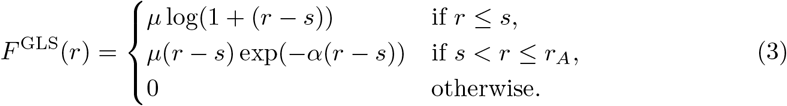

In all three functions *µ* denotes the spring stiffness (split between repulsive and adhesive interactions for the piecewise quadratic force). Furthermore, *r*_*R*_ denotes the interaction distance for repulsive interactions for the piecewise quadratic force and *α* controls the width of the exponential decay of the GLS force in the adhesive regime (see [24] for an in-depth discussion and other force function examples). These one-dimensional forces are extended to two or three dimensions by multiplication with the normalized direction vector between cell mid-points, i.e. we define the pairwise force vector between cells *i* and *j* as 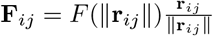, where **r**_*ij*_ = **x**^(*j*)^ − **x**^(*i*)^ and **x**^(*i*)^ and **x**^(*j*)^ denote the midpoint coordinates of cells *i* and *j*. Other force functions implemented in CBMOS include the linear force [29, 30] and the Hertz force [31, 32].

The cells are assumed to move in a microenvironment with a very low Reynolds number [33] in which inertial effects can be neglected. Under this assumption, the update equation for the midpoint coordinates of the *i*th cell is

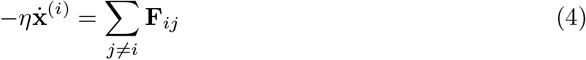

where the drag force proportional to the cell’s velocity is balanced with the force being exerted on the cell by its neighbors. The drag coefficient *η* acts as a scaling of the pairwise interaction forces and can thus be arbitrarily fixed as *η* = 1.

Given an initial placement for the coordinates of the population, Equation (4) is solved numerically at discrete time points. Methods implemented in CBMOS for doing this include the explicit first-order forward Euler method [34] (commonly used by other center-based model simulation software such as Chaste [14] and MecaGen [15]), the explicit second-order Midpoint [35] and Adams-Bashforth [34] methods (the latter is used by PhysiCell [16]), as well as the implicit first-order backward Euler method [34]. We refer to Section 3.2 for a detailed description and comparison of the forward and backward Euler methods.

If necessary, the numerical solving of Equation (4) is interrupted by proliferation events taking place. CBMOS implements cell division by choosing a random cell division direction and placing two daughter cells of equal radius *R* at a fixed distance in that direction such that the former position of the mother cell is located in the center. By default, CBMOS uses an initial separation distance between daughter cells of 0.3 cell diameters.

### 2.2 Design overview

The mathematical model described in the previous section relies on several basic modeling assumptions, most prominently the force functions describing pairwise interactions between cells. Furthermore, it includes many numerical aspects such as the method used to solve the ODE system. Choosing all model components and their parameters — such as cutoff radii and spring stiffness values for the force functions or time step sizes for the numerical solver — as well as understanding their effects on the accuracy and efficiency of the simulation are open problems. To our knowledge, the majority of center-based model software are targeted at modelers addressing biological problems and the numerical components mentioned above are usually not exposed to the user. These software are therefore ill-suited for investigating numerical questions. CBMOS is designed to fill this gap.

CBMOS allows to probe the interplay of different model components — mainly the pairwise interaction force and the numerical solver — and their combined effect on the population level behavior as well as on the efficiency of the simulation. To do so, CBMOS provides a flexible, easy-to-use interface that is easily expandable to study the effects of force functions, ODE solvers, time step sizes or cellular events in the context of center-based models. CBMOS implements a number of pairwise force functions found in the literature and other popular software packages for the simulation of center-based models, as well as five ODE solvers, including three second-order solvers and one implicit solver. A simple example showing how to set up and run a simulation of two cells can be seen in Listing 1, with a more complex example being described on CBMOS’ documentation webpage^5^. Furthermore, the interested user can find code examples on common numerical analysis workflow scenarios in the Jupyter notebooks available in the GitHub repository (see the examples folder, as well as the code belonging specifically to this and our previous publication [24]).

The CBMOS code is event-driven, meaning that cell events are queued according to their execution time and the mechanical equations for the center positions are solved in between the execution of cell events, see Figure 1 for the general structure of the code. The event-driven implementation is advantageous for the numerical study of center-based models as it avoids an additional splitting error that arises when simulations proceed in a step-driven manner. Nevertheless, we allow for cellular events to be aggregated at fixed time points to improve simulation efficiency when a large number of cellular events need to be handled, as is commonly done in major modeling software. Currently, CBMOS only supports cell divisions as cellular events.

**Figure 1:**
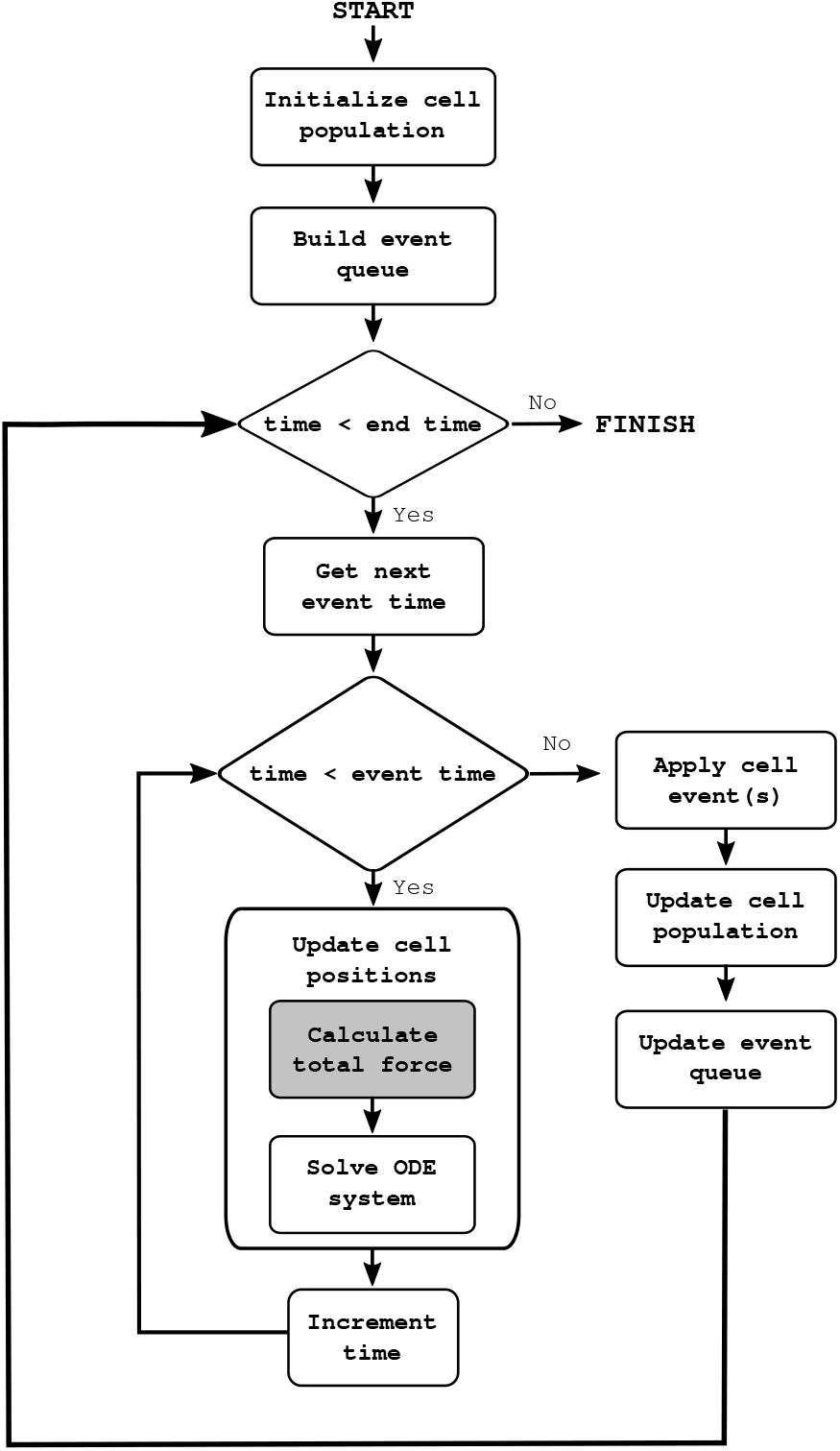
Simplified flow chart for the CBMOS code. The positions of the cell midpoints are calculated numerically with the ODE solver between any two consecutive cell events. At that point, the next event is resolved and new events are queued. This process is repeated until the end time is reached. If the aggregation of cellular events is needed for efficiency, the event queue is built with event times rounded to the next possible event time according to the desired resolution. The main bottleneck consists of the calculation of the total force, highlighted with a darker background. This is where CuPy is used as a drop-in replacement for NumPy when access to a GPU is available.

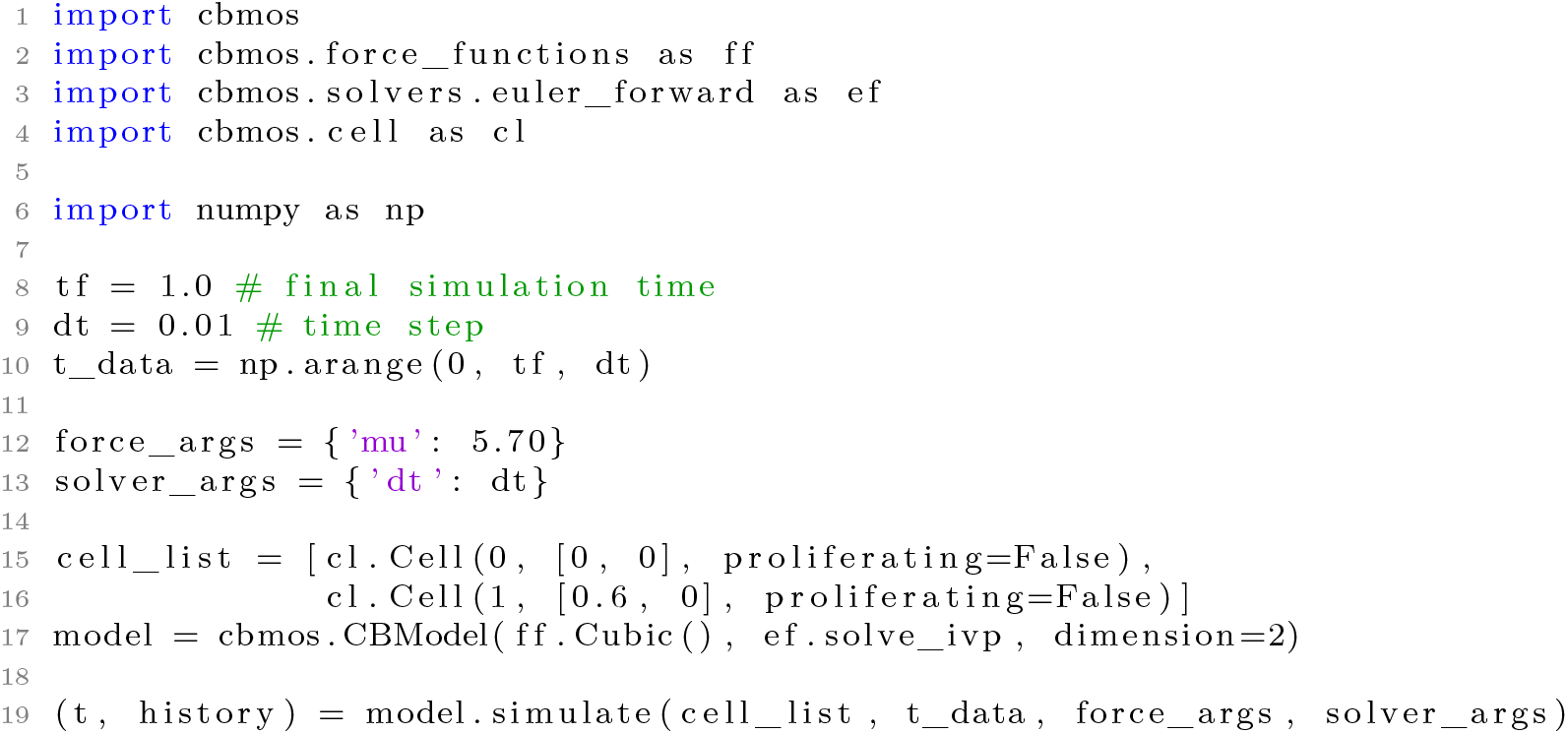

Listing 1: Minimal working example showing how to initialize and simulate a cell population in CBMOS. First of all, a cell list is created, specifying the initial position of each cell, as well as if and how they divide. Secondly, a model is initialized with the desired force function, numerical solver, and dimension (options are one, two or three space dimensions). As a pairwise force, a model can take any function that takes a floating point number (the distance between two cells) as well as any number of keyword arguments, and returns another floating point number (the resulting force). Likewise, the model can take any numerical solver that implements the same interface as SciPy’s ODE solver `scipy.integrate.solve_ivp`. Finally, parameters for the force functions, the solver — typically the time step, or threshold values for implicit solvers — and the times at which the position of the cells should be recorded are passed to the `simulate` function of the model itself. The latter returns the full history of the cell population as a list of cells for each time step, as well as the time points themselves. The `simulate` function also takes a number of optional parameters — for instance, to fix the seed of the random number generator and ensure the simulations are reproducible — all of which are described in the documentation.

### 2.3 Implementation

Simple center-based model implementations typically target cell counts from a few hundred to a few thousand cells. Larger system sizes are possible by using parallel implementations in a compiled language as done e.g. by PhysiCell [16], Chaste [36] or Biocellion [20]. Additionally, the ya‖a framework [17] achieves high performance by executing CUDA/C++ code on GPUs. We follow a complementary approach in our code as its main purpose lies in facilitating the numerical study of the model components. As such, performance is secondary to fast prototyping, but nevertheless the possibility of simulating more cells in less time would allow for more realistic test problems and is thus very desirable. The three main points of our approach can be summarized as (i) the use of the Python programming language to enable fast prototyping [37], (ii) the use of array programming via NumPy to achieve a reasonable performance on the CPU and (iii) the use of a high-level GPU-library as a replacement to NumPy which enables the transfer of the most computationally expensive portions of the code to the GPU. This speeds up the simulations by a factor of up to 30 for our typical test problems on a modern and widely accessible GPU, allowing for larger system sizes while still retaining the advantage of having a fast and easily accessible development cycle.

The computational bottleneck of a center-based model implementation is the calculation of the total force acting on each cell. According to this force the cell position is updated in every time step in Equation (4). In order to calculate the total force vector for the complete cell system, all pairwise forces need to be evaluated. In other major center-based software, computing the force interactions between cell pairs is usually done using a bounding box technique, where space is discretized into voxels larger than the maximum interacting distance between cells [38]. The forces applied to a given cell are then only computed for the cells located in the same voxel and in neighboring voxels. Given that the system usually relaxes to a given density of cells per voxel, i.e. the number of cells per voxel will be bounded from above, this algorithm achieves linear complexity with respect to the number of cells. Implementing such an algorithm in pure Python code, however, is typically orders of magnitude slower than in compiled languages.

The naive implementation which calculates the interactions between all possible cell pairs, on the other hand, scales quadratically with the number of cells. However, it is easily expressed with array programming, a programming paradigm based on elementary array operations, such as indexing, vectorization, broadcasting and reduction. In Python, NumPy is now the *de facto* standard for array programming [39]. By using an optimized, pre-compiled layer of C code under the hood, NumPy provides improved performance for all array operations, while still making it possible to write legible Python programs. On top of that, there exist a myriad of Python modules implementing NumPy’s array protocol. Such libraries include Dask [40], for distributed large arrays or PyData/Sparse [41] for sparse matrices. For GPU computations in particular, there exist several high-level libraries aiming to extend NumPy, such as CuPy [26], MinPy [42] (deprecated, now merged with MXNet Gluon [43]) and afNumPy [44]. In practice, such array implementations provide drop-in replacements for a large subset of NumPy functions and thus require only minimal modifications of the code to be used.

If a GPU is available, a CBMOS user can specify the use of the CuPy library as a high performance computing backend. Although originally developed specifically for 3D graphics, GPUs have become widely available for general computations in recent years with the advent of the CUDA programming language and GPGPUs (general-purpose computation on GPUs) [45]. When threads are relatively independent and only have to synchronize for atomic operations, GPUs make it possible to run massively parallel applications executing thousands of threads at a time. In the case of CBMOS, the bulk of the computations is done when computing the total force which can be expressed as independent, predictable operations applying to all elements of an array, making it well-suited for GPU computations. The arrays involved in the computation of the total force are created on the GPU and calculation of the force vector is done there. This GPU-enabled version reduces the computation time for a single evaluation of the force vector for 10,000 cells to 0.4 seconds, which is about thirty times faster than on the CPU.

In the numerical experiments in Section 3.1, we compare our array-based approach both on the CPU and the GPU to the bounding box algorithm outlined above.

### 2.4 Code availability and usage

The full program is readily available under the MIT license on the Python package index^6^ and on GitHub^7^. Installing and running CBMOS is straightforward as it only depends on a few well maintained external modules (mainly the Python scientific software stack NumPy, SciPy and matplotlib and optionally CuPy). The documentation is available on the project’s GitHub page^8^ and describes how to set up a simple simulation. An example of a convergence study is also presented there.

One of the main advantages of CBMOS over other similar software is the possibility to run simulations, analyse them and interpret the results all in a single Jupyter Notebook. Jupyter Notebooks have become very popular in recent years and are an excellent way to report reproducible scientific findings [37]. In fact, the recent development of online platforms providing free, ready-to-use resources to execute such notebooks (even on GPUs) makes this process even easier. For instance, all the notebooks used in this study are freely available on our GitHub repository and can be set up and rerun in a couple minutes on Google Colab [27, 28].

## 3 Numerical experiments

We now proceed to demonstrate how CBMOS can be used to perform numerical experiments. The following section has two main parts. In Section 3.1, we focus on the computational performance of CBMOS both on CPU and GPU and draw practical bounds as to which one is most suitable, depending on the number of cells considered in the simulation. In Section 3.2, to show-case what kind of questions can be addressed using CBMOS, we conduct a numerical study comparing the implicit backward Euler method for solving the update equation to the more commonly used explicit forward Euler method. Note that while all numerical experiments were done in two dimensions, CBMOS is capable of simulating three dimensional cell populations as well.

### 3.1 Performance comparison CPU/GPU

In this section, we illustrate the performance gain enabled by evaluating the total force vector on the GPU instead of the CPU, pushing the limit of how many cells a center-based code written in Python can simulate. First, we ran a performance benchmark to study wall time as a function of the number of cells for a compressed monolayer relaxing to steady state. Second, we prescribed a fixed wall time and counted how many cells could be simulated in a monolayer growth experiment within that time. For both experiments we compared our array-programming-based implementation to the bounding box algorithm described in Section 2.3. Taken together, these experiments provide a practical estimate for the most suitable algorithm in terms of cell population sizes.

The benchmarks were run on Snowy, an HPC cluster provided by the Multidisciplinary Center for Advanced Computational Science (UPPMAX). The node we used consisted of two 8-core Xeon E5-2660 processors at 2.2 GHz, 128 GB of memory, and was equipped with an Nvidia T4 GPU.

#### Relaxation benchmark

In the first benchmark scenario we generated cell populations of different sizes arranged in a compressed honeycomb pattern in which the distance between any two neighboring cells was initialized to 0.8 cell diameters (the rest length was set to *s* = 1.0 cell diameter). We then allowed the system to relax to steady state over the course of one in-simulation hour. Within this time no proliferation took place, so that no cellular events needed to be handled.

The force function was chosen as the generalized linear spring (GLS) force with parameter settings *µ* = 1.95 and *a* = − 2 log(0.002*/µ*). The choice of these parameters resulted in a relaxation time between daughter cells after cell division of one hour (in-simulation time), as described in [24]. The time step was set to Δ*t* = 0.1*h*, ensuring that cell trajectories after cell division remain physically correct (again, for details we refer to our previous numerical study [24]).

Figure 2a shows the total execution time for the relaxation experiment with the three implementations described in Section 2.3: (i) the bounding box implementation (denoted by ‘Box’ in the legend), (ii) the array implementation using NumPy and running only on the CPU (denoted by ‘NumPy’) and (iii) the array implementation using CuPy in addition to NumPy. The latter transfers the calculation of the force vector at each time step to the GPU and is denoted by ‘CuPy’ in the legend. Although the bounding box implementation has the best computational complexity, it was an order of magnitude slower than both array implementations for all practical use cases considered here (as a reference, in the previous CBMOS publication [24], we only considered experiments with 2, 38, 74 and 400 cells). The bounding box implementation only beat the NumPy implementation starting at 10^4^ cells and above, at which point the simulation took around five minutes to complete.

**Figure 2:**
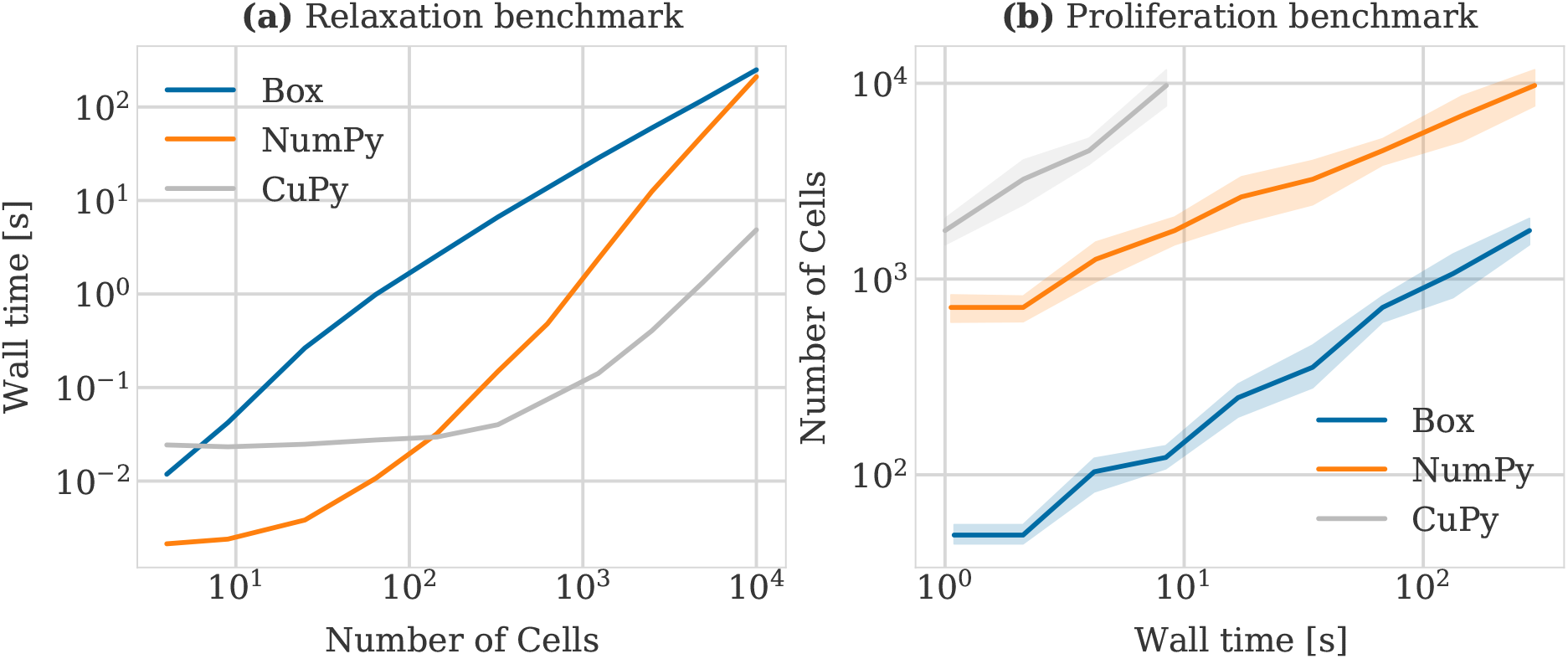
(a) Wall time as a function of the number of cells in a relaxation experiment. Cells started in a compressed honeycomb shape where distances were reduced by 20% of the rest length and the simulation was run until the system relaxes. (b) Number of cells simulated as a function of the wall time measured in seconds. The simulation started with a single cell that proliferated according to an exponentially distributed cell cycle duration. At regular time intervals, the simulation was stopped and the amount of cells was counted. Each simulation was run five times with a different seeds, from which we drew the 95% confidence intervals. A time step of 0.1 hour and the GLS force with *µ* = 1.95 and *a* = − 2 log(0.002*/µ*) were used for all simulations in both benchmarks. For the bounding box algorithm, the box size was equal to the cutoff distance, namely 1.5 cell diameters.

Transferring the calculation of the force vector — the main computational bottleneck — to the GPU brought significant improvements starting already from 100 cells. More specifically, the GPU-enhanced array implementation was up to about one and a half orders of magnitude faster for the range of 100 to 10,000 cells. For lower numbers of cells its performance was dominated by the overhead of transferring the data to and from the GPU. Most notably, for 10^4^ cells, the simulation only took 4 seconds, compared to around five minutes for the bounding box implementation and the CPU-only array implementation. Above this point, the GPU we used ran out of memory, although theoretically in terms of pure execution times (given enough memory), it would be faster than the bounding box implementation up to 6.25 *×* 10^5^ cells.

Overall, this benchmark illustrates the range of application of the CBMOS package in terms of cell population sizes for non-proliferating two dimensional populations and in the context of exploratory prototyping, where single realizations should run within a minute. For up to 10^2^ cells, the NumPy implementation is sufficient. Above this threshold, the use of a GPU enables significant performance gains for the simulation of up to 10^4^ cells, decreasing the simulation time from several minutes to just a few seconds. For larger system sizes, the memory requirements become prohibitive and other center-based software frameworks should be considered.

#### Monolayer growth benchmark

In a second benchmark, we studied the question of how many cells one could simulate with the different versions within a fixed execution time in a monolayer growth experiment. To this end, we set up a single initial cell. This initial ancestor proliferated according to an exponentially distributed cell cycle duration (with a mean of 1.0 hour), generating a large cell population over time. We stopped the simulation after a fixed wall time had elapsed and counted the number of the cells in the population. We considered simulation times of up to a few hundred seconds, which in our opinion is about the maximum reasonable waiting time one can afford to wait when prototyping. Division events were only allowed to take place every 0.1*h*. All other parameters, in terms of force functions and time steps, were the same as in the relaxation benchmark.

Figure 2b shows our results by plotting the number of cells simulated as a function of wall time. Again, in spite of its quadratic complexity, the NumPy implementation outperforms the bounding box implementation by an order of magnitude in terms of the number of simulated cells for all practical simulation times. Deploying the code on a GPU showed even greater performance, simulating around 10,000 cells in about 8 seconds, at which point the GPU ran out of memory.

In the relaxation benchmark the number of cells was fixed from the beginning and no proliferation took place, meaning no cell events needed to be handled during the simulation. This allowed the solver to run continuously from the beginning to the end. In this benchmark, however, proliferation was included and division events took place throughout the simulation, usually forcing the solver to restart after every time step once the cell population had grown past a certain size. This benchmark shows that even in this case the CuPy implementation largely outperforms the two other options.

Having shown that our implementation is reasonably efficient for system sizes of up to 10,000 cells (when using the GPU) in different experimental settings, we now move on to study the use of an implicit method for solving the system of ODEs for the cell midpoints.

### 3.2 Numerical study of the implicit backward Euler method

In this section we show-case the ability of CBMOS to study the numerical properties of center-based models. In particular, we investigate whether the use of an implicit method for solving the system of ODEs governing the movement of the cell midpoints over time is beneficial in terms of computational cost necessary to achieve a desired accuracy.

#### Comparison of the stability of the forward and backward Euler methods

A commonly used numerical method for solving the ODE system for the center positions in center-based model implementations is the forward Euler method [34]. It is an explicit method, meaning the function value at the next time point can be explicitly calculated from the current function value. For an initial value problem stated as

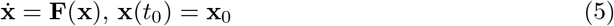

the forward Euler method can be written as

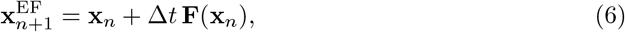

where **x**_*n*_ denotes the current function value, **x**_*n*+1_ the function value at the next time point *t*_*n*+1_ = *t*_*n*_ + Δ*t* and Δ*t* is the step size. As an explicit method, the forward Euler method suffers from stability issues. More specifically, if the time step size Δ*t* is chosen too large, the numerical solution will oscillate and grow without bounds, even though the true solution does not. The backward Euler method – the simplest implicit method for solving ODE systems – does not exhibit this constraint on the time step size [46]. It can be written as

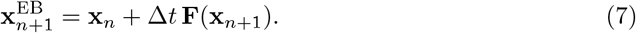

Note that here, in contrast to Equation (6), the total force **F** (which in general is non-linear) is evaluated at the next time point. Thus, obtaining a value for the next function value **x**_*n*+1_ with the backward Euler method requires the iterative solving of Equation (7) with a linear system solve in each step.

In this section, we illustrate the different stability properties of the forward and backward Euler methods using the simple test case of two cells relaxing after cell division. The cells were initially placed at a overlap of 0.3 cell diameters which they eliminated until they were at rest at a distance of one cell diameter with no forces acting between them. Parameters have been chosen such that the duration of this process — called the relaxation time — was (arbitrarily) fixed as one hour. As shown in our previous study [24], using the forward Euler method requires the time step size to be chosen lower than a certain stability threshold to recover a numerically stable solution. Moreover, if cell trajectories after division should be physically correct, i.e. cells should not jump apart before adhering again, then the time step size is even more restricted by a monotonicity bound which is half the stability bound for this specific test case. The exact values of these bounds are force function and parameter dependent (see [24] for details).

Figure 3 shows the behavior of the numerical solution for the cell trajectories after cell division when calculated using the forward (left column) and backward (right column) Euler methods. Subsequent rows differ in the size of the time step used, ranging from a very small time step size of Δ*t* = 0.025 *h* in panels (a) and (b), to a large step size of Δ*t* = 0.125 *h* in panels (e) and (f). All panels show the trajectories for three different pairwise force function choices, illustrating that stability is both a property of the solver and the ODE system itself (defined via the pairwise force function). For reference, the dotted curves in each panel correspond to an accurate solution (less than 1% relative error) calculated using Δ*t* = 0.005*h*. Note that the first two panels from the left column were regenerated using the same data as in our previous publication [24]. In the left column, where the explicit forward Euler method was used, we observe that for the smallest time step size all trajectories are physically correct, whereas for the larger step sizes the trajectories show physically correct (piecewise polynomial force in (c) and GLS force in (c) and (e)), physically incorrect yet stable (cubic force in (c) and piecewise polynomial force in (e)) or even numerically instable behavior (cubic force in (e)), depending on how sensitive the force functions are to the time step choice.

**Figure 3:**
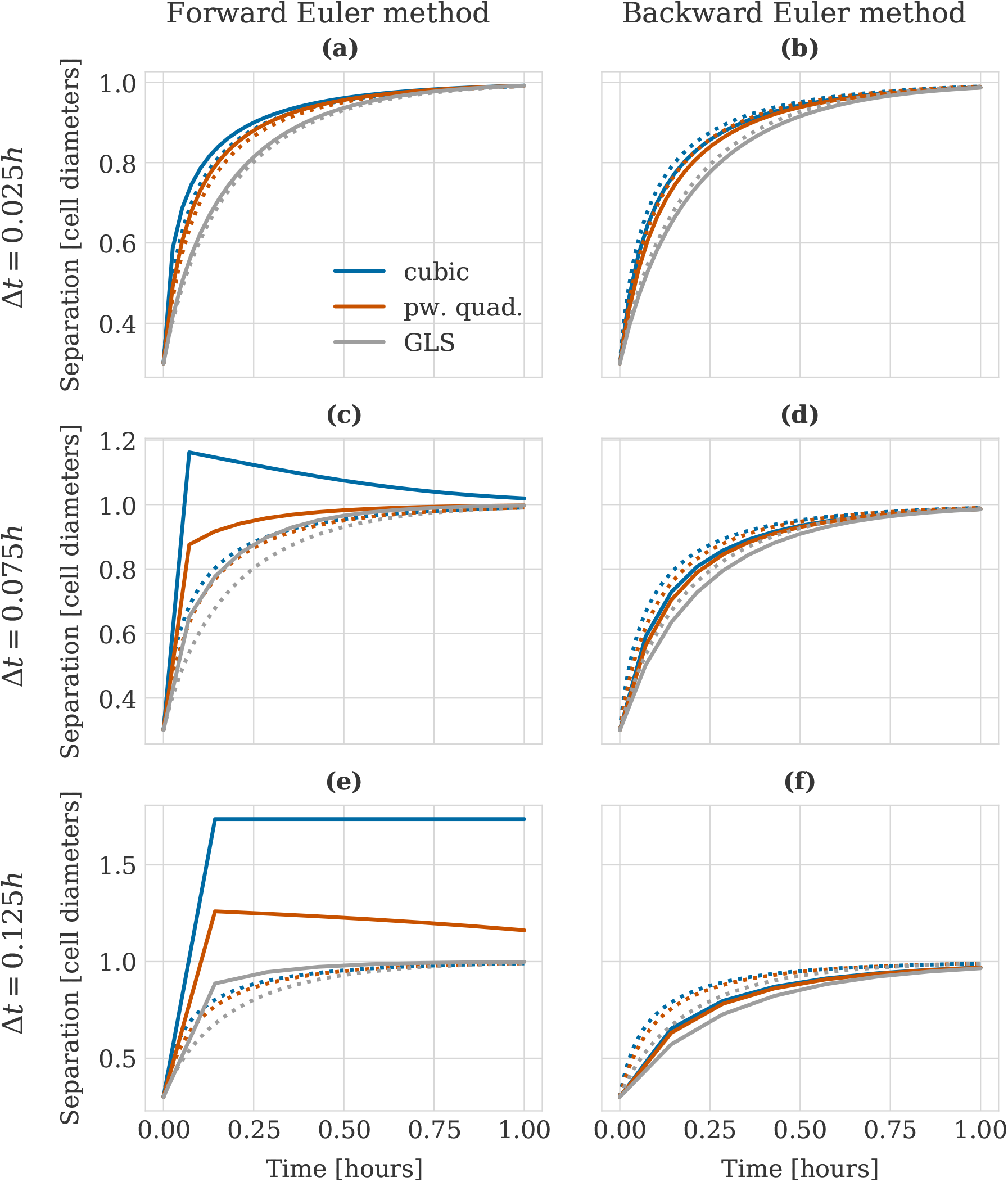
Stability of the numerical solution for the relaxation dynamics between daughter cells under different force functions when calculated using the forward Euler method (left column) and the backward Euler method (right column). The different rows use successively larger time step sizes: (a, b) Δ*t* = 0.025*h*, (c, d) Δ*t* = 0.075*h*, (e, f) Δ*t* = 0.125*h*. The legend shown in panel (a) is valid for all panels. For reference, the dotted curves correspond to an accurate solution (less than 1% relative error) calculated using Δ*t* = 0.005*h*. Parameters for the force functions were chosen as *s* = 1.0 cell diameters, *r*_*A*_ = 1.5 cell diameters, *µ*_cubic_ = 5.7, *µ*_*R*_ = 9.1, *µ*_*A*_ = 1.911, *r*_*R*_ = 1.18029, *µ*_GLS_ = 1.95, *α* = 7.51. The first two panels of the left column of this figure were regenerated using data from [24].

In contrast, the right column of Figure 3 shows the exact same experimental setup for the implicit backward Euler method. Here, we observe that independently of the time step size and the force function choice the trajectories remain physically correct. (Note of course that for larger time step sizes the accuracy decreases as expected). Our further investigations in [24] illustrated that a choice of a too large time step when using the forward Euler method — in particular a time step size violating the monotonicity bound at the pairwise cell level — could result in geometrical differences even at the population level. It is therefore of interest to resolve the pairwise dynamics correctly and hence the question arises whether the backward Euler method is a more suitable choice because it allows for larger time step sizes without stability nor monotonicity constraints.

Before we turn to this question of computational efficiency, we conduct a convergence study, the results of which are described in the next section.

#### Convergence study of the forward and backward Euler methods

Aside from stability, an important property of a numerical method is its order of convergence, describing how fast its accuracy increases when decreasing the time step size. It can be shown theoretically that both the forward and backward Euler methods are first-order methods, meaning that roughly speaking halving the step size also halves the error in the numerical solution. To confirm this numerically, we conducted a convergence study where we analysed the error as a function of the time step size for both methods and different example systems. We measured the error relatively to a reference solution **x**_ref_ generated with a very small time step of Δ*t*_ref_ = 10^−4^ *h* as

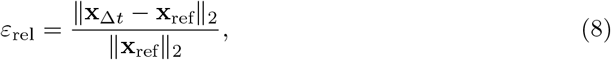

where the coarser solution **x**_Δ*t*_ was interpolated down to the fine time grid used for the generation of **x**_ref_.

Figure 4 shows the results of our convergence study for four different experimental setups. The first setup, shown in panels (a) and (b), considered the relaxation between two daughter cells after cell division, same as before when comparing the stability properties in the previous section. In the second, shown in panels (c) and (d), we placed two cells at an initial distance of 1.15 cell diameters and let them adhere until they touched. The third setup in panels (e) and (f) was chosen as a small monolayer population of initially 19 cells. All cells were allowed to divide at the beginning of the simulation, leading to a highly compressed cell population which then relaxed to a steady state configuration over the course of two in-simulation hours. Finally, as a fourth experimental setup we considered the relaxation of a large monolayer, similarly to the setup of the relaxation benchmark in Section 3.1. Here, we initialized a monolayer population of 400 cells for which we were able to conduct the convergence study in a reasonable time by using CuPy as a backend. The results for this experimental setup are shown in panels (g) and (h).

**Figure 4:**
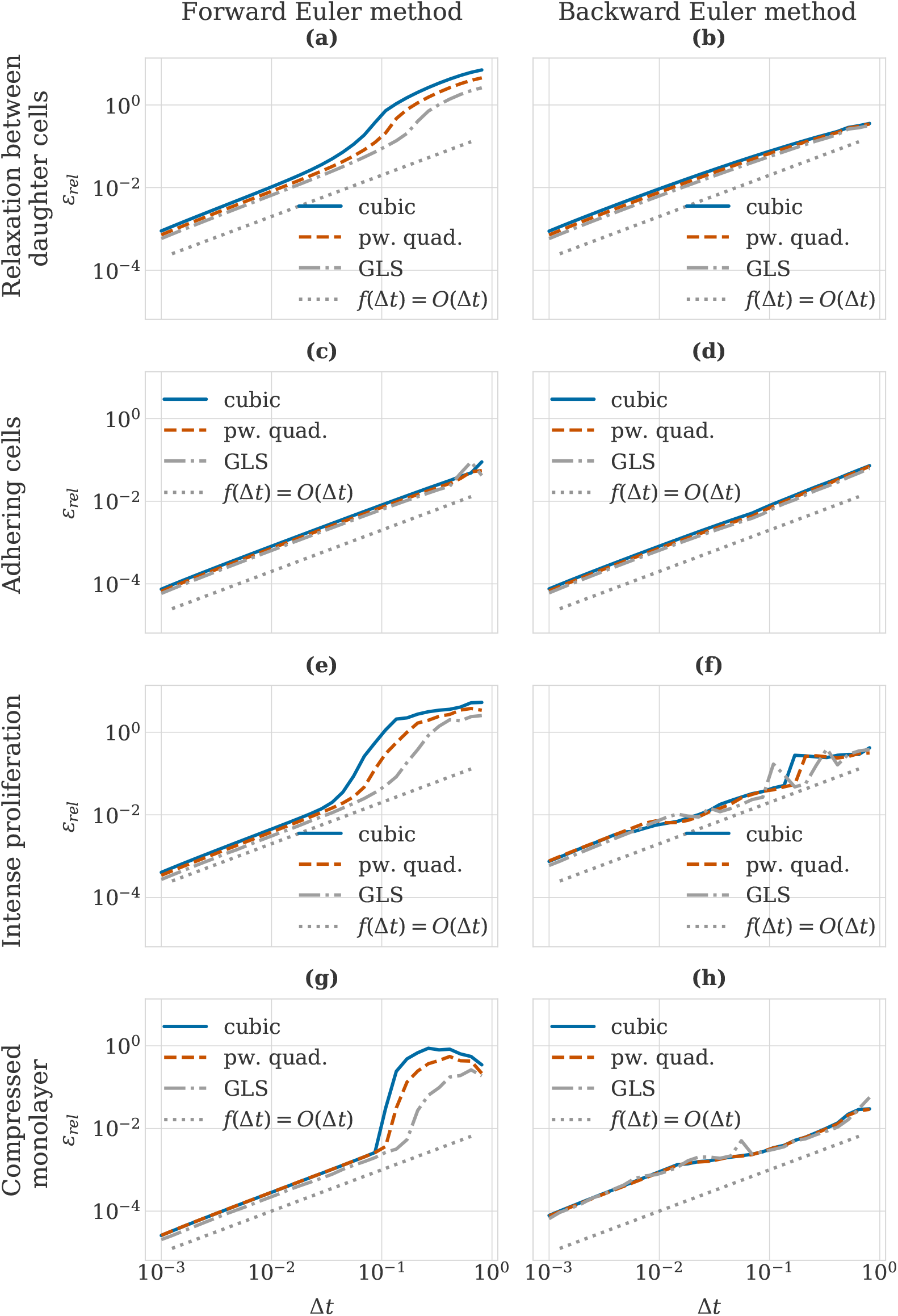
Impact of time step size on the relative error for the different combinations of forces and both numerical solvers tested on four model problems (i)-(iv). (a-b) Convergence study for the relaxation experiment (i), initial separation 0.3 cell diameters, (c-d) convergence study for adhering cells (ii), initial separation 1.15 cell diameters, (e-f) convergence study for a monolayer population of initially 19 cells under intense proliferation (iii) and (g-h) convergence study for a compressed monolayer population of 400 cells (iv). The numerical solver used was in (a, c, e, g) the forward Euler method, and in (b, d, f, h) the backward Euler method. The axes of the plots are in logarithmic scale. The dotted lines show a linear function to facilitate observation of the convergence order. Note that the first three panels in the left column include data used also in our previous publication [24].

In the left column of Figure 4 the forward Euler method was used to calculate cell trajectories, whereas the right column shows the results for the backward Euler method. All plots show the relative error as a function of time step size for three different force functions with the axes in logarithmic scale. We observe that both numerical methods show the correct first-order convergence across all force functions and experimental setups for small time step sizes Δ*t <* 10^−2^ *h*. For larger time steps in the range of 0.05 *h* to 0.5 *h* the forward Euler method shows a larger relative error for all experimental setups except the second one with the adhering cells. This is due to the loss of monotonicity and stability in the cell trajectories for these time step sizes, as discussed in the previous section. This means that the backward Euler method is able to achieve relative error values of the order of 10^−1^ with larger time step sizes than the forward Euler method. This raises the question whether the backward Euler method can be beneficial if cell trajectories need to be resolved with only moderate accuracy, i.e. when a relative error of 5 − 10% is acceptable.

In the next section we consider the additional cost of using the backward Euler method and measure its execution time in comparison to the forward Euler method in order to answer the question whether it is actually worth it to use the backward Euler method or if using the forward Euler method with small time steps is more efficient.

#### Additional cost of the backward Euler method

Implicit methods such as the backward Euler method gain their improved stability properties at an increased computational cost. As they formulate the equation for the next function value depending on the gradient at the next time point (instead of the current as explicit methods do), they require this equation to be solved iteratively with a linear system solve in each step. The function value **x**_*n*+1_ in Equation (7) is calculated by solving

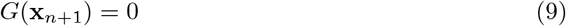

by Newton iterations, where *G*(**x**) = **x − x**_*n*_ − Δ*t***F**(**x**). More specifically,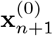 is initialized using **x**_*n*_ and then for several iterations *j* the following two steps are executed,

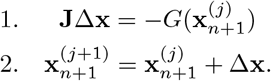

In the first step 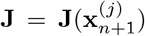 denotes the Jacobian of G which directly depends on the Jacobian of the ODE system 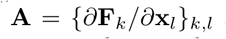 as **J** = **I −** Δ*t***A**, where **I** is the identity matrix. **J** defines the linear system that needs to be solved for Δ**x**. This can be done e.g. with the generalized minimal residual method (GMRES) [47]. In our implementation, we stop the Newton iterations once the relative difference between iterates is smaller than a threshold *ϵ* _Newton_ = min(10^−3^, Δ*t*). Similarly, we pass `atol=tol`=min(10^−3^, Δ*t*) to the GMRES implementation provided by the `scipy.sparse.linalg` module [48]. Choosing smaller threshold values increases the accuracy at an increased computational cost due to more iterations. The values we use were chosen as the largest values that still ensured the correct order of convergence for the backward Euler method in our numerical experiments.

CBMOS provides an analytically correct implementation of the Jacobian **J**. In practice, however, it is more efficient to approximate the matrix-vector product **Jv** in GMRES via

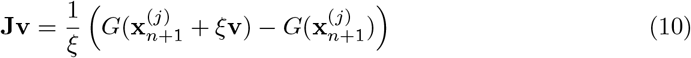

without having to assemble the Jacobian **J** [49]. Here, *ξ* denotes an approximation parameter, for which we use *ξ* = 0.001 in all our numerical experiments.

In order to evaluate whether using the backward Euler method can be more efficient for moderate accuracy values, we measured the wall time for the convergence study executed in the previous section. More specifically, we averaged wall times over 10 repetitions of calculating the cell trajectories. Figure 5 shows this average as a function of the relative error for the four different experimental setups. We note that for moderate accuracy requirements, the backward Euler method is able to use larger time step sizes than the forward Euler method. Nevertheless, using the backward Euler method is about one order of magnitude slower than using the forward Euler method across the complete range of relative error values for both experimental setups with two cells and the intense proliferation test case. It is only for the case of the large compressed monolayer and a relative error of roughly 3% or larger that the backward Euler method achieves a similar wall time to the forward Euler method. At this point the backward Euler method was able to use time step sizes up to nearly 8 times as large depending on the force function chosen. The exact values for relative errors and time step sizes are compared in Table 1.

**Table 1:** Table showing the relative error values *ε*_*rel*_ and the time step values Δ*t* (in hours) for the forward and backward Euler methods in combination with different force functions. The experimental setup chosen was the relaxation of a compressed monolayer of 400 cells. Values were rounded to three decimals.

|  | Forward Euler |  | Backward Euler |  |
| --- | --- | --- | --- | --- |
| | $\varepsilon_{rel}$ | $\Delta t$ | $\varepsilon_{rel}$ | $\Delta t$ |
| Cubic force | 0.031 | 0.108 $h$ | 0.030 | 0.808 $h$ |
| Pw. quad. force | 0.031 | 0.136 $h$ | 0.029 | 0.808 $h$ |
| GLS force | 0.027 | 0.212 $h$ | 0.030 | 0.646 $h$ |

**Figure 5:**
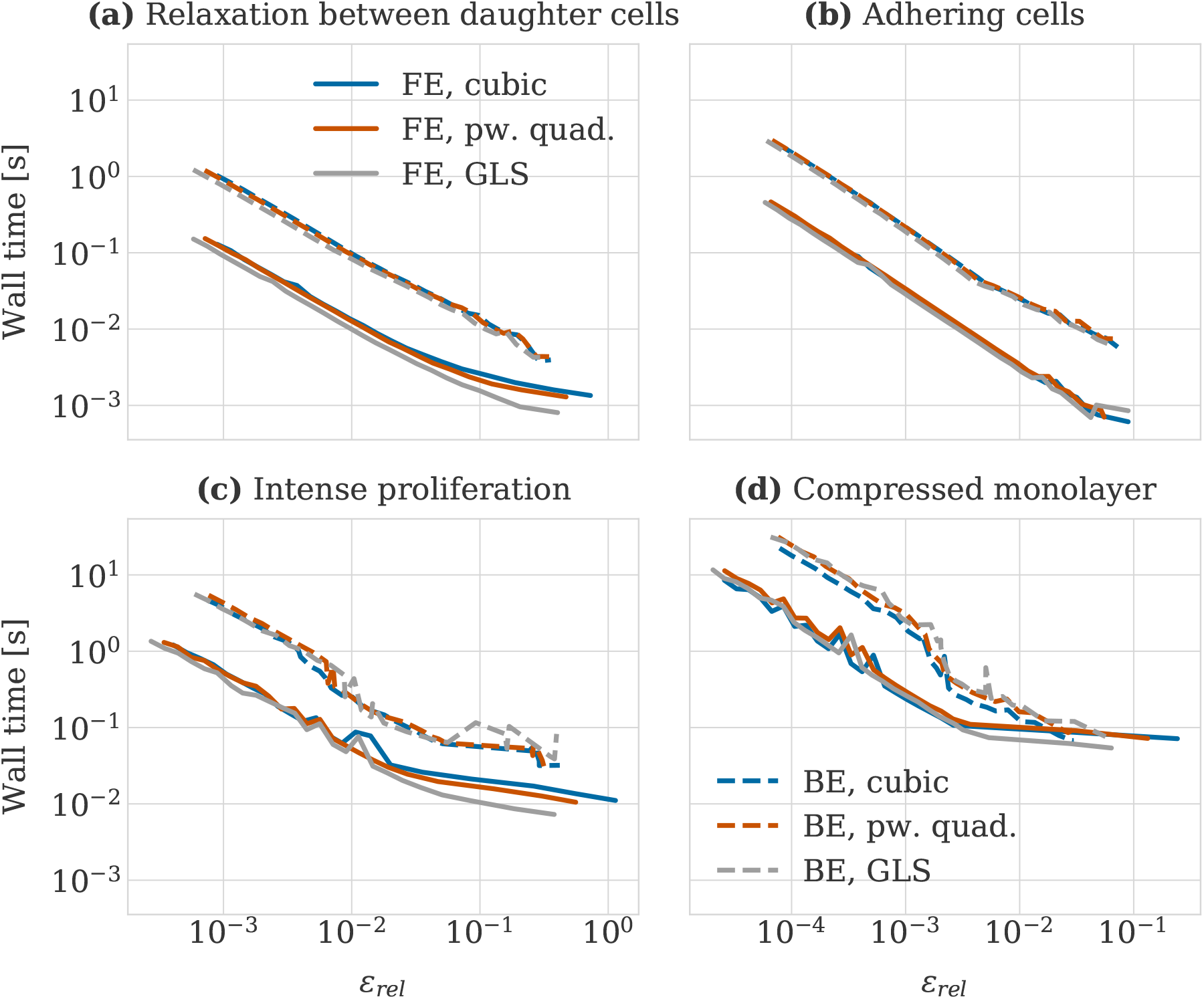
Average wall times for calculating cell trajectories as a function of relative error with respect to a reference solution for different force function (cubic, piecewise quadratic and GLS) and solver (forward and backward Euler) combinations. The panels show the results for the four different experimental setups considered: (a) relaxation between daughter cells, (b) adhering cells, (c) small monolayer under intense proliferation and (d) compressed monolayer of 400 cells (simulated using CuPy as a backend). In each panel full lines represent experiments using the forward Euler (FE) method, whereas for the dashed lines the backward Euler (BE) method was used (the legends in panels (a) and (d) are both valid for all panels).

Implicit methods, such as the backward Euler method, can be beneficial when stability, rather than accuracy, is the limiting factor [34]. In this subsection, we have investigated if this was indeed the case for ODE systems from center-based models. We conclude that based on our numerical experiments it is not beneficial to use the backward Euler method compared to the forward Euler method, even if it allows for larger time step sizes for low accuracy values. Its additional computational cost due to requiring a linear solve remains too high even when approximating the Jacobian and using as few iterations as possible.

## 4 Discussion

In this article we presented CBMOS, an open-source package for the numerical study of center-based models. CBMOS provides a flexible interface to study the effect of core components of the numerical simulation of center-based models — amongst others the force function used for the pairwise interaction forces between cells and the numerical method for solving the update equation for the cell midpoint coordinates — on the mechanics of a proliferating cell population as well as on the efficiency of the simulation itself. To this end, it includes implementations of many popular force functions as well as several explicit and implicit first- and second-order numerical methods. Its interface is designed to allow for easy extension in terms of more force functions or numerical methods, but also more types of cellular events.

Written in Python, CBMOS provides easy access for modelers of any level and enables fast prototyping workflows. Thanks to relying on NumPy’s array programming paradigm it is reasonably efficient simulating small cell populations on the order of hundreds of cells on the CPU. Moreover, it extends this range to cell populations of up to 10,000 cells in connection with a high-end GPU through the CuPy library. In that case execution times for simulating monolayer growth are on the order of a few seconds, with the amount of memory available on the specific GPU becoming the limiting factor. In its current state on our test hardware CBMOS was capable of running repeated convergence studies on test systems with thousands of cells within a few hours.

Naturally, an implementation in a compiled language (as used for other open source simulation packages for center-based models such as e.g. PhysiCell [16] or Chaste [14]) or accessing CUDA features more directly (as done e.g. in ya‖a [17]) can be expected to be more performant in terms of pure execution speed. However, those kinds of implementations usually require a substantial investment in terms of both development and implementation time, not to mention that they may require a steep learning curve for users less familiar with progamming. It is our hope that developing a Python package will have the additional benefit of being more easily accessible for modelers not coming from a computational background. In general, the combination of a interpreted language with high-level GPU-libraries such as CuPy can be very attractive and advantageous as the former permits fast-prototyping workflows, while the latter ensures significant performance gains at virtually no added implementation time cost.

CBMOS is implemented in an event-driven fashion, meaning it simulates the mechanics of the cell population until the next cellular event happens. This has the advantage that cellular events are applied at the exact time they occur. For large populations with many cellular events this approach can become inefficient due to the need to simulate the mechanics for fractions of a time step in order to advance the system to the correct state before applying the cellular event. Therefore, we have chosen to augment CBMOS with the option to aggregate cellular events to a fixed time resolution. This means that at pre-determined time points we apply all cellular events that would have taken place between the last check point and the current time. This time-driven approach is how most center-based model frameworks are implemented. It results in a lower bound to the step size of the mechanics simulation at the cost of an additional splitting error. Thanks to implementing both approaches CBMOS could be used in future work to investigate the error of the time-driven versus the event-driven approach. This would allow to formulate guidelines on how to choose the fixed time resolution of the cellular events in order to balance incurred error with simulation efficiency.

In a previous publication [24] we used a prototype of CBMOS to investigate how popular force functions should be parametrized to reproduce consistent mechanical behavior for two dimensional cell populations, as well as how the combination of force function and numerical solver affects the efficiency of center-based models in terms of time step sizes. For the latter we focused on explicit first and second-order solvers. In this article, to further illustrate the types of questions CBMOS can be used to address, we studied the complementary question of whether an implicit method for solving the update equations can be beneficial. More specifically, we studied whether the better stability properties of the backward Euler method can balance out its increased computational cost compared to the forward Euler method for different experimental setups. In line with our previous study we considered three popular force function choices on the test cases of (i) daughter cells relaxing after cell division, (ii) adhering cells and (iii) a small monolayer population undergoing intense proliferation. Furthermore, we considered a large compressed monolayer of 400 cells as an additional setup, the simulation of which was possible thanks to using CuPy as a GPU-backend. For all experimental setups the backward Euler method was less efficient in terms of wall time for computing the cell trajectories than the forward Euler method, although it was able to use larger time step sizes for moderate to small accuracy values. On average, the backward Euler method was around one order of magnitude slower than the forward Euler method, even with generous threshold settings and when approximating the Jacobian at the cost of an extra evaluation of the total force (instead of assembling the complete analytical Jacobian at a higher cost). These results extend K. Atwell’s findings in her thesis [23], where she investigated the backward Euler method as well as another second-order implicit method for a small tumor-growth experiment.

To summarize, our results confirm that using the forward Euler method with sufficiently small time step sizes is computationally more efficient than using the backward Euler method, at least as long as fixed time-stepping is used. With adaptive time stepping, where the time step size is chosen dynamically according to a suitable error estimate, there might be gains with the backward Euler method for systems that spend long durations in states where overall the forces between cells are weak, e.g. when strong compression forces due to cell division events are rare. Additionally, in such a setting it might be even more advantageous to switch between both methods, depending on whether the time step size is restricted by accuracy or stability concerns. We leave the exploration of both ideas for future work.

There exist several other center-based model frameworks implementing a larger feature set and better performance due to being written in a compiled language. Nevertheless, we believe that, by making it possible to isolate fundamental aspects of the simulation through a user-friendly API, CBMOS offers easy access to any modeler wishing to quickly challenge key numerical properties of their center-based model or to test preliminary hypotheses before committing to a more complex simulation framework.

## Acknowledgments

We would like to thank Per Lötstedt for fruitful discussions about the implicit Euler method, its implementation and how to measure its cost. We would also like to thank Carl Nettelblad for providing a singularity container with CuPy installed. All numerical experiments were performed on the Snowy compute resources provided through the Uppsala Multidisciplinary Centre for Advanced Computational Science (UPPMAX) within the Project SNIC 2019-8-227. This work has received funding from the Swedish Research Council under grant 2015-03964 and from the eSSENCE strategic initiatives on eScience.

## Footnotes

1 https://github.com/somathias/cbmos

2 https://github.com/somathias/cbmos

3 https://somathias.github.io/cbmos/

4 https://research.google.com/colaboratory/

5 Available at https://somathias.github.io/cbmos/

6 https://pypi.org/project/cbmos/

7 https://github.com/somathias/cbmos/

8 https://somathias.github.io/cbmos/

